## Supplementary material for "Generic comparison of lumen nucleation and fusion in epithelial organoids with and without hydrostatic pressure": Supp_Note_Theory

### Supplementary Information Theory:

#### 1 Purpose of theoretical modeling

In this Supplementary Note, we address theoretical questions for lumen nucleation and fusion dynamics in epithelial organoids. As mentioned in the main text, we study three different systems with varying initial cell numbers. Experimental parameters such as cell proliferation rate, cell-cell adhesion, cortical tension, luminal pressure potentially vary across the three systems. If the biochemical dynamical processes are considered, difference in subtlety may diverge. Nevertheless, three systems have common features such as initial lumen nucleation phase and succeeding lumen fusion phase followed by a single lumen state surrounded by cell layers. There also remains discrepancies between MDCK/pancreas and epiblast. In the former systems, lumen nucleates between two cells, while lumen nucleates only after 10 cells merge into rosette structures in epiblast. Why does this commonality and individuality emerge?

We propose answering these questions through theoretical modeling and simulations. The hypothesis is that these processes are mechanically controlled, i.e. (1) mechanical force balance between lumen and cell layers determines the condition for the lumen growth or shrink, and (2) lumen fusion process is controlled by the Young-Laplace type pressure driven dynamics, then (3) the final single lumen state is dynamically balanced state of cell proliferation and lumen growth, finally (4) difference in dynamics of epiblast from MDCK/pancreas is mainly caused by low luminal pressure in epiblast.

In order to validate these hypotheses, we introduced a multi-cellular phase field model (PFM) (Section 2 of this Supplementary Note) and simplified analytical argument based on a nucleation theory (Section 3 of this Supplementary Note). Both models share a common free energy. The former model allows us to simulate detailed shapes of cells and lumens even with irregular deformations of lumen and cells as in real experiment. This is the advantage of PFM compared with such as the vertex model. Cell rearrangement such as T1 process can be automatically achieved in PFM, while in the usual vertex model, reconnections between vertices have to be made in T1 process. Comparison with other computational models were summarized in our paper[1] . We do not assume any symmetries for each cell shape and positions except for the initial conditions. Once the initial conditions are set, the model equations spontaneously evolve, cells grow and divide, and lumens nucleate if the mechanical conditions are met. If the resultant dynamics such as lumen number, cell number, fusion dynamics, and final single lumen state are statistically similar, we believe that the real systems also follow the same principle. To have mathematical insight, employing a simplified model system is useful, because it enables analytical prediction for the condition of lumen nucleation. The second model assumes symmetry among cells, such as equal shape and volume for each cell with spherical lumen and spherical outer shape of organoids to allow the stability analysis of the systems.

Assumptions of the models are as follows. Shape and volume of each cell evolves in the direction to minimize associated energy. Development of the lumen also satisfies the steepest descent dynamics of the associated energy. ECM elasticity is also considered. The growth of the whole system is driven by three factors: (1) each cell tends to grow toward its target volume. (2) If the cell volume exceeds the predetermined threshold volume, cell can divide as far as the time duration after previous division exceeds the minimum division time which mimics the duration of cell cycle. (3) Each lumen grows or shrinks depending on the balance between the driving force for the lumen growth (one can compare this driving force parameter as the osmotic pressure. See Sec.1.8) and tension from the surrounding cell layers. (We proved that in our PFM, lumen volume evolves according to the well-known equation, Eq. (21), for the lumen dynamics when the lumen is spherical.)

Surface tension, cell-cell adhesion, cell-ECM adhesion, cell volume elasticity are included in the energy terms. All parameters in the PFM are non-dimensionalized. If it requires, quantitative comparison between the model parameters and physical unit is possible in PFM as demonstrated in A. Badillo[2]. However, in the present work we compare the relative strength of cell cortical tension, cell-cell adhesion, and lumen osmotic pressure for different types of cysts, since all coefficients are non-dimensionalized. This limits the comparison between simulation and experiment within qualitative level. In the analytical calculation of the postulated free energy of the cysts, we chose surface tension of cells and luminal hydrostatic pressure close to the values reported in experiments. Therefore, the comparison is semi-quantitative for the analytical model.

### 2 Multicellular phase-field model for lumen fusion simulations

#### 2.1 Multi-Cellular Phase Field Model

In this Supplementary Note, we describe the mathematical model used to simulate the organoid growth dynamics. Important physical assumptions here are as follows: (i) All cells have volume regulation, cell-cell adhesion, cortical tension, and excluded volume effect and every cell has the same property. (ii) The driving forces of lumen growth (osmotic pressure) are the same and constant to time for all lumens included within each organoid. The hydrostatic pressure of the lumen is automatically determined by the balance with the tension of the cell layer in this model. To implement this, we apply the multi-cellular phase field method developed by Nonomura (2012)[3] and Akiyama et al 2018)[4].

Organoid dynamics are simplified into three components: cell, lumen, and ECM or surrounding medium. Geometries of these components are represented by the corresponding virtual fields called phase fields in the following. $u_m(\boldsymbol{r},t)(m=1,\dots,M)$, $\mathrm{s}(\boldsymbol{r},t)$, and $\mathrm{c}(\boldsymbol{r},t)$ are the phase field variables representing the shapes of each cell, lumen, and ECM (medium) respectively. Here cell index is denoted by $m$ with $M$ the total cell number. The regions of each phase field with $1$ and $0$ correspond to inside and outside of each component.

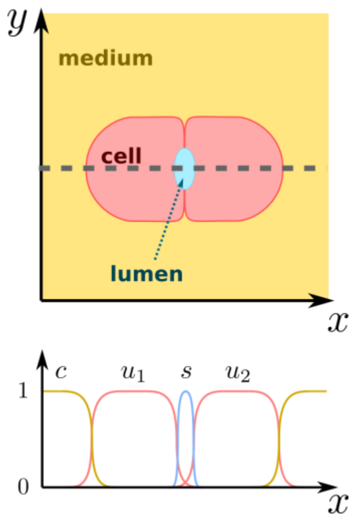


Figure S1: *Schematic representation of the model. (Top) Phase field model of cells, lumens, and ECM or medium in 2D. (Bottom) Cross-sectional view of the phase fields along the dotted line. Red, cyan, and yellow colors correspond to the phase fields corresponding to cells, lumen, and ECM or medium, respectively, for both the curves (top) and the regions (bottom).*

#### 2.2 Governing equations for dynamics of phase fields

The dynamics of the fields is based on minimization of the free energy through gradient descent. We used the construction principle based on Akiyama et al (2018)[4]. In addition, we newly take into account the existence of ECM, which provides mechanical resistance for organoids to grow and, therefore, can influence the steady states. In addition, cell growth control (2.3), cell cortical tension control (2.4), and deciphering the role of osmotic pressure (2.5) are newly included. The total free energy of the system is defined as,

$$E = E_u + E_s + E_c + E_{int}, \qquad (1)$$

where $E_u, E_s, E_c$ are the energy for cell, lumen, ECM, respectively. $E_{int}$ is the interaction energy.

Each term is further subdivided into terms caused by volume change, surface energy, osmotic pressure, and so on.

$$\begin{aligned} E_u &= E_{pf}(u) + E_{vol}(u) + E_{surf}(u), \\ E_s &= E_{pf}(s) + E_{osm}(s), \\ E_c &= E_{pf}(c) + E_{vol}(c) + E_{osm}(c), \\ E_{int} &= E_{adh} + E_{excl}\,, \end{aligned} \qquad (2)$$

where $E_{pf}$ is phase field model specific energy terms defined by $E_{pf}(u) = \sum_m \int \left[\frac{D_u}{2}|\nabla u_m|^2 + \frac{1}{4}u_m^2(1 - u_m)^2\right] d\boldsymbol{r}$, $E_{pf}(s) = \int \left[\frac{D_s}{2}|\nabla s|^2 + \frac{1}{4}s^2(1-s)^2\right] d\boldsymbol{r}$, and $E_{pf}(c) = \int \left[\frac{D_c}{2}|\nabla c|^2 + \frac{1}{4}c^2(1-c)^2\right] d\boldsymbol{r}$ for the cell variable u, lumen variable s, and ECM variable c, respectively. The coefficients $D_u, D_s,$ and $D_c$ are positive constants representing diffusion coefficient of $u$, $s$, and $c$. These double-well potentials are defined in such a way that each variable takes on the values 0 or 1, with an interface in between.

$E_{vol}$ terms are the excess energy terms due to volume change from its natural volume, and defined as $E_{vol}(u) = \sum_m \frac{\alpha_u}{12}(V - \int h(u_m)d\boldsymbol{r})^2$ for cell, and $E_{vol}(c) = \frac{\alpha_c}{12}(V_c - \int h(c)d\boldsymbol{r})^2$ for ECM. $\alpha_u$ and $\alpha_c$ are positive constants representing bulk modulus of cells and bulk modulus of ECM, respectively. $V$ and $V_c$ are target volume of cells and target volume of ECM respectively and $h(x) \equiv x^2(3 - 2x)$.

$E_{surf} = \sum_m \frac{\gamma}{12}\int |\nabla h(u_m)|^2 \, d\boldsymbol{r}$ represents the surface energy of cells, with $\gamma$ the surface tension of the cell. $E_{osm}(s) = -\frac{\xi}{6}V_l = -\frac{\xi}{6}\int h(s)d\boldsymbol{r}$ is the energy coming from osmotic pressure, defined by the multiple of the osmotic pressure of the lumen $\xi$ and the lumen volume $V_l$ defined as $V_l = \int h(s)d\boldsymbol{r}$. $E_{osm}(c) = -\frac{\xi_c}{6}\int h(c)d\boldsymbol{r}$ is the energy that comes from the osmotic pressure of ECM. $\int h(c)d\boldsymbol{r}$ is the volume of ECM.

$E_{adh}$ is adhesion energy defined by

$$E_{adh} = \sum_m^M \sum_{m' \neq m}^M \frac{\eta_u}{12}\int \nabla h(u_m) \cdot \nabla h(u_{m'})d\boldsymbol{r} + \sum_m^M \frac{\eta_{cu}}{6}\int \nabla h(c) \cdot \nabla h(u_m)d\boldsymbol{r}, \tag{3}$$

Where $\eta_u$ is the cell-cell adhesion energy, $\eta_{cu}$ is the cell-cell adhesion energy per area (length in 2D).

$E_{excl}$ is the volume exclusion term defined by

$$E_{excl} = \sum_m \left[\sum_{m' \neq m} \frac{\beta_u}{12}\int h(u_m)h(u_{m'})d\boldsymbol{r} + \frac{\beta_{su}}{6}\int h(s)h(u_m)d\boldsymbol{r} + \frac{\beta_{cu}}{6}\int h(c)h(u_m)d\boldsymbol{r}\right] + \frac{\beta_{cs}}{6}\int h(c)h(s)d\boldsymbol{r},$$
(4)

Where $\beta_u, \beta_{su}, \beta_{cu},$ and $\beta_{cs}$ are the coefficient of the excluded volume interaction between cells, that between cells and lumens, between ECM and cells, and that between ECM and lumens. The ECM is assumed to have pressure and elasticity, controlled by $\xi_c$ and $\alpha_c$, respectively. The ECM also has volume exclusion interactions with the cells and the lumen, respectively, and has adhesion with cells.

If each variable evolves by gradient descent of the free energy

$$\tau_u \frac{\partial u_m}{\partial t} = -\frac{\delta E}{\delta u_m}, \tau_s \frac{\partial s}{\partial t} = -\frac{\delta E}{\delta s}, \tau_c \frac{\partial c}{\partial t} = -\frac{\delta E}{\delta c}, \tag{5}$$

the time evolution equations for each variable are obtained as follows, although we adopted different time evolution for the cell cortical tension control (see Sec. 1.3):

$$\tau_u \frac{\partial u_m}{\partial t} = D_u \nabla^2 u_m + u_m(1-u_m)\left(u_m - \frac{1}{2} + f_u\right), \tag{6}$$

$$f_u = \alpha_u \left(V_m(t) - \int h(u_m)dr\right) + \gamma_u \nabla^2 h(u_m)$$

$$-\beta_u(\psi_u - h(u_m)) - \beta_{su} h(s) - \beta_{cu} h(c),$$

$$+\eta_u \nabla^2(\psi_u - h(u_m)) + \eta_{cu} \nabla^2 h(c), \tag{7}$$

$$\tau_s \frac{\partial s}{\partial t} = D_s \nabla^2 s + s(1-s)\left(s - \frac{1}{2} + f_s\right), \tag{8}$$

$$f_s = -\beta_{su}\psi_u - \beta_{cs} h(c) + \xi, \tag{9}$$

$$\tau_c \frac{\partial c}{\partial t} = D_s \nabla^2 c + c(1-c)\left(c - \frac{1}{2} + f_c\right), \tag{10}$$

$$f_c = \alpha_c \left(V_c - \int h(c)dr\right) - \beta_{cu}\psi_u - \beta_{cs} h(s) + \eta_{cu} \nabla^2 \psi_u + \xi_c, \tag{11}$$

where $\tau_u$, $\tau_s$, and $\tau_c$ are time constants for evolution. Here we set $\tau_u = \tau_s = \tau_c = 1.0$.

##### 2.3 Cell growth control

When $\alpha_u$ is a constant value, the growth rate of cell volume is determined by the ratio of the time constants $\tau_u/\tau_s$. However, if $\tau_u$ is changed, terms other than the volume growth rate (surface tension and cell-cell adhesion terms) are also affected, preventing accurate control volume alone. Therefore, we control the time scale of cell growth by introducing the following time evolution of the target volume $V_m(t)$ and changing the time scale of the rate of increase of the target volume.

$$\tau_V \frac{dV_m(t)}{dt} = \bar{V} - V_m(t) \tag{12}$$

where $\tau_V$ and $\bar{V}$ are positive constants. By replacing $V$ in the $E_{vol}(u)$ term with $V_m(t)$, the volume of each cell grows following the target volume $V_m(t)$ that is based upon a time constant $\tau_V$. Thus, the time scale of cell growth can be controlled by $\tau_V$.

#### 2.4 Cell cortical tension control

In the phase-field method using the Allen-Cahn form, the phase-field interface has an intrinsic surface tension due to the diffusion constant and double-well potential terms. Since changing the coefficients in these terms also changes the interface width, it is not possible to control the interface width and surface tension independently. Therefore, in this simulation, we used the method of Olsson(2005)[5] and Badillo(2012)[2] to eliminate the above intrinsic surface tension and keep the interface width constant for all phase field variables: cell, lumen, and medium. We consider the application of the resharpening method (Olsson(2005)、Badillo(2012)[2,5] to the phase field variable $\phi$ in the following time evolution equation,

$$\tau\frac{\partial\phi}{\partial t} = D\nabla^2\phi + \phi(1-\phi)\left(\phi - \frac{1}{2} + f\right) . \tag{13}$$

Instead of integrating Eq. (13), it can be separated into two steps,

$$\tau\frac{\partial\phi}{\partial t} = \phi(1-\phi)f , \tag{14}$$

$$\frac{\partial\phi}{\partial t^*} = \nabla\cdot\left(D\nabla\phi - \sqrt{2D}\phi(1-\phi)\frac{\nabla\phi}{|\nabla\phi|}\right). \tag{15}$$

The first equation gives the phase change, and the second equation is used to control the interface profile. After the phase $\phi$ is integrated by Eq. (14) for a time step of $\Delta t$, $\phi$ is allowed to evolve in an auxiliary time $t^*$ according to the conservative Eq. (15) until it converges ($\frac{\partial\phi}{\partial t^*} = 0$) to satisfy hyperbolic tangent profile, $\phi = \frac{\left[1+\tanh\left(\frac{r}{\sqrt{2D}}\right)\right]}{2}$. In this study, the resharpening method is applied to Eqs. (6), (8), and (10) to control the interface width and independent surface tension which is determined by $\gamma$ in the $E_{surf}$ term.

#### 2.5 Computational implementation

Dynamics of the phase-field variables is calculated by performing the following two operations alternately: [I] Genuine time evolution— the set of the phase-field variables (all cells $u_m$, lumen $s$, ECM $c$) is updated for only a single step of time $\Delta t$ by transforming Eq.(6),(8), and(10) as in the transformation from Eq.(13) into Eq.(14); [II] Resharpening— the steady state of each phase field (every cell $u_m$, lumen $s$, ECM $c$) is computed based on Eq. (15). The parameter values used in the main text are: the size of simulation area $\Omega = 20.48 \times 20.48$, a spatial grid size $\Delta x = 0.01$, a time step $\Delta t = 0.02, D_u = D_s = D_c = 0.001, \alpha_u = 1.0, \alpha_c = 0.001, \beta_u = \beta_{su} = \beta_{cu} = \beta_{cs} = 1.0, \eta_u = 0.0075, \eta_{cu} = 0.001, \gamma = 0.0057, V = 3.0, V_c = \Omega$.

In lumen fusion simulations for 8 cells' initial cell number condition (Fig. 6b), each cell divides into two daughter cells when the cell volume condition $V_d = 2.9$ is satisfied. The definition of the cell division plane was based on Akiyama et al. (2018). A small lumen area of the radius $r_s = 0.5$ creates on the center of division plane. If the driving force $\xi$ of the lumen is greater than the pressure from the surroundings, the lumen will grow, but if not, it will disappear naturally by the dynamics. In case of circular lumen, this condition is consistent with Laplace's law.

In the experiment configured by attaching two epiblast cysts, relatively fast fusion dynamics due to active cell motion was observed. To mimic this experiment, we performed lumen fusion simulation with two cysts. In lumen fusion simulation with two cysts (Fig.8b-f), cells do not divide, and we introduced two model modifications that are "Active force of cells" and "Volume conservation of lumens".

Applying the volume conservation of lumens in this two cysts simulation can be validated when the fusion process is enough fast compared to lumen growth/decay dynamics governed by Eq. (21) or Eq. (25).

[I]Active force of cells— In simulations for epiblast fusion, active forces were introduced to consider the cell motion observed in experiments of epiblasts (Fig. 4i, j, k). The m-th cell is subjected to the active force $\boldsymbol{F}_m(t)$defined as:

$$\boldsymbol{F}_m(t) \equiv \mathrm{sgn}(r_m(t) - r_{th})\frac{\boldsymbol{r}_m(t)}{|\boldsymbol{r}_m(t)|}, \tag{16}$$

where $\boldsymbol{r}_m(t) = \boldsymbol{G}_0(t) - \boldsymbol{G}_m(t)$, $\boldsymbol{G}_0(t)$ is the center of mass of the two cysts, $\boldsymbol{G}_m(t)$ is the center of mass of the *m*-th cell, $r_m(t) = |\boldsymbol{r}_m(t)|$, $r_{th}$ is a given threshold distance, and

$$\mathrm{sgn}(x) = \begin{cases} 1\ (x > 0) \\ 0\ (x = 0) \\ -1\ (x < 0) \end{cases}. \tag{17}$$

here we set $r_{th} = 3.0$.

Gm is driven by the force defined by Eq. (16) and (17) as follows:

$$\tau_g \frac{\partial \boldsymbol{G}_m(t)}{\partial t} = \boldsymbol{F}_m(t) \tag{18}$$

where $\tau_g$ is a time constant of evolution. Here we set $\tau_g = 500$.

[II] *Volume conservation of lumens*— To conserve lumen volume, we introduce new terms for volume conservation same that of cells in the free energy $E_s$. The lumens in the two cysts are represented by separate phase field variables $s_1(\boldsymbol{r}, t)$ and $s_2(\boldsymbol{r}, t)$, respectively. The volume conservation term for lumens is defined as follows:

$$E_{s_i} = \frac{\alpha_s}{12}\left(V_{s_i}(t) - v_{s_i}\right)^2 (i = 1,2) \tag{19}$$

where $\alpha_s$ is a coefficient of balk modulus of lumen, $V_{s_i}(t)$ is a target volume of $s_i$, and $v_{s_i}(t) = \int h(s_i) d\boldsymbol{r}$.

$V_{s_i}(t)$ evolves in time as in Eq. (12) and increases to $\bar{V}_{s_i}$ that the target volume of $V_{s_i}(t)$. In these simulations, we set $V_{s_i}(0) = v_{s_i}(0)$, $\bar{V}_{s_i} = 3v_{s_i}(0)$, and $\alpha_s = 1$.

##### 2.6 Implication of the driving force for the lumen growth

In the following argument, we relate ξ in the dynamics of the lumen variable to the osmotic pressure.

In the luminogenesis, the lumen growth dynamics is often expressed as

$$\frac{dV_l}{dt} = \lambda_w A_{apical}(\Pi - \Delta p), \tag{20}$$

where $\lambda_w$ is the water permeability, $A_{apical}$ is the area of apical side of cell layer surrounding the lumen, $\Pi$ is the osmotic pressure, $\Delta p$ is the hydrostatic pressure difference across the cell layer. We demonstrate that lumen growth dynamics in PFM obey the similar equation that enable us to give the implication of $\xi$ in Eq. (9). For this purpose, it is useful to separate the free energy related to the lumen variable $s$ as

$$E'_s = E_{pf}(s) - \frac{\xi}{6}V_l + E_{int}, \tag{21}$$

In one dimensional (1D) case, such as the 1D cut in Fig.S1, the integral in $E_{pf}(s)$ can be easily evaluated, while the $E_{int}$ term originated from the excluded volume interaction vanishes in 1D case. Because cell's position can slide without restoring force as the lumen region expands in 1D. By using he analytic solution of the interface in 1D, $s(x) = (1/2)\tanh\,(x/(2\sqrt{2D}))$, the first term reads

$$E_{pf}(s) = \int \left[\frac{D_s}{2}|\nabla s|^2 + \frac{1}{4}s^2(1-s)^2\right] d\boldsymbol{r} = \frac{\sqrt{2D_s}}{12}. \tag{22}$$

In two dimension (2D), when the lumen is circular with a radius R (thus $V_l \sim \pi R^2$) and assuming that the interface is thin enough compared to $R$, $E_{pf}(s)$ for circular lumen becomes $\frac{\sqrt{2D}}{12}$ times circumference length $2\pi R$; i.e. $E_{pf}(s) = (\sqrt{2D}/12)(2\pi R)$. Therefore, the dynamics of $V_l$ obeys

$$\frac{\partial V_l}{\partial t} = -k\frac{\partial E'_s}{\partial V_l} = k\left(\frac{\xi}{6} - \frac{\partial E_{pf}(s)}{\partial R}\frac{\partial R}{\partial V_l}\right) = k\left(\frac{\xi}{6} - \frac{\sqrt{D_s/2}}{6R}\right), \tag{23}$$

where $k$ is the kinetic coefficient and it is inversely proportional to $\tau_s$ in Eq. (5). This equation can be compared to Eq. (21) with the Laplace law like relation, $\Delta p = \sqrt{D_s/2}\,/(6\mathrm{R})$, where $\sqrt{D_s/2}\,/6$ is the tension of the lumen surface. $\Pi = \xi/6$. In 2D, cell layer surrounding the lumen adds additional tension due to the exclusive interaction between lumen variable and cells through the term, $E_{int} = \sum_m \frac{\beta_{su}}{6}\int h(s)h(u_m)d\boldsymbol{r}$ by interacting with cell variable $u_m$. Each $u_m$ is further interacting with other cells and ECM by adhesion and exclusion, thus as the lumen volume increases cell layer will add additional tension. The total force from the cell layer to the lumen is difficult to calculate analytically, however, if the tension $t$ of the cell layer is uniform along the circular apical surface, $E_{int} \sim 2\pi R t$ and $\frac{\partial R}{\partial V_l} = \frac{1}{2\pi R}$ hold in 2D, and $E_{int} \sim 4\pi R^2 t$ and $\frac{\partial R}{\partial V_l} = \frac{1}{4\pi R^2}$ hold in 3D. Finally, we obtain

$$\frac{\partial V_l}{\partial t} = -k \frac{\partial E_s'}{\partial V_l} = k\left(\frac{\xi}{6} - \frac{\partial E_{pf}(s)}{\partial R}\frac{\partial R}{\partial V_l} - \frac{\partial E_{int}(s)}{\partial R}\frac{\partial R}{\partial V_l}\right) = k\left(\frac{\xi}{6} - \frac{\sqrt{2D_s}/6 + t}{R}\right), \quad (24)$$

for 2D case. By denoting the effective tension, $T = \sqrt{2D_s}/6 + t$, the last term can be compared to the Laplace law, $\Delta p = T/R$. In 3D case, $\Delta p = 2T/R$ holds. After performing the cell cortical tension control described in Sec.1.3, $\sqrt{2D_s}/6$ term vanishes. The tension $T$ of the cell layer may depend on thickness of the cell layer and the elasticity of ECM as well. Therefore, $T$ is not a constant but generally depends on $R$ in nonlinear manner. This $R$ dependence of $T$ gives rise to the saturation of lumen growth. Although real value of $T$ is difficult to evaluate analytically, we can estimate it from the simulation dynamics which gives how $\frac{\partial V_l}{\partial t}$ and $R$ grow in time for fixed value of $\xi$. From the fitting of the time series of $\frac{\partial V_l}{\partial t}$ and $R$ by using Eq. (25), we found an empirical relation in our model that $\Delta p$ is smaller than $\xi$ but proportional to $\xi$ over the range of $R$ values tested in simulation.

#### 2.7 Code availability

The software code used for the simulation is available in the Github, https://github.com/kana-fuji/MCPFM_for_Lumen_Fusion.git

### 3 Free Energy of Organoid and Condition for Lumen Formation

We consider the free energy of the organoid consisting of N cells with a single lumen embedded in ECM,

$$F = -\Delta p V_A + \gamma_A A_A + \gamma_L \sum_{i=1}^{n} A_L^i + k \sum_{j=1}^{N} \left(V_c^j - V_c^0\right)^2 + k_E (V_E - V_E^0)^2 \quad (25)$$

where $\Delta p$ is the pressure difference across the lumen and ECM, $V_A$ is the volume of the lumen, $\gamma_A$ and $\gamma_L$ are surface tension of cell apical and lateral area, respectively. $A_A$ is the total apical area, $A_L^i$ is the i-th cell lateral area, $n$ is the number of lateral surfaces, $k$ is the elastic constants for cell volume change, $V_c^j$ is the volume of j-th cell, $N$ is the number of cells, $V_c^0$ is the natural volume of the cell, $k_E$ is the elastic constant of ECM, $V_E$ is the volume of ECM, and $V_E^0$ is the natural volume of the ECM. We consider N=2, 4, 8 cases by assuming spherical lumens, spherical outer shape for the organoid and ECM surrounding it, equal shape and equal size for each cell, for simplicity. Thus, $A_L^i$ ($V_c^j$) takes the same value for all i (j). For N=2 case, cell shape is assumed to be a hemisphere, For N=4, each cell is a quadrant sphere obtained by cutting sphere with the equatorial plane and the plane passing 0°and 180° meridians. For N=8 case, each quadrant sphere is cut into two to make 8-segment spheres. In each case, the center of the organoid is occupied by a single spherical lumen. Therefore, each hemisphere is missing a center with a self-similar hemispherical lumen for n=2 case. The same for N=4 and 8 cases. Therefore, the number of lateral surfaces becomes n=2, 8, and 24 for N=2,4, and 8 case, respectively. Denoting the lumen diameter r and organoid diameter R, lumen volume $V_A = \frac{4\pi}{3} r^3$, apical area $A_A = 4\pi r^2$ are common for N=2.4, and 8 cases. Lateral surface term, $\gamma_L \sum_{i=1}^{n} A_L^i$ becomes $\frac{2n}{N}\pi(R^2 - r^2)\gamma_L$ for N cell case. Cell volume contribution, $k \sum_{j=1}^{N} \left(V_c^j - V_c^0\right)^2$ becomes $\frac{1}{N} k g_s^2 (R^3 - r^3 - R_0^3)^2$ for N cell case, where $g_s = \frac{4\pi}{3}$ and $V_c^0 \equiv \frac{1}{N} g_s R_0^3$ is the natural volume of each cell. ECM elasticity term $k_E(V_E - V_E^0)^2 = k_E g_s^2 (L^3 - R^3 - L_0^3)^2$ is common for all cases, where L is the system size, $g_s L_0^3$ is the natural volume of the ECM. We minimize the free energy F with respect to r and R for different values of $\Delta p$. For the visualization, we plot $F(r, R_\min)$using the value of $R = R_{min}$, where $F(r)$ is minimized with respect to $R$ for each $r$ in Fig. 6f-h.

Condition for the Lumen formation is determined by the balance between the tendency to minimize total surface area under the influence of cell and ECM volume elastic energy. As shown in Fig.1a, when $\Delta p = 0$ [Pa], F is minimum at r=0 (no lumen) for N=2 and 4 cases implying stable rosette, r=4.4 um for N=8 case, and r=12.3 um for N=12 case, implying lumen can be formed only when number of cells is larger than 8. In this case, organoid has a rosette like shape without lumen for N=4 case. The situation is similar at $\Delta p = 30$ [Pa] in Fig.1b. At higher pressure $\Delta p = 100$ [Pa], lumen can be created for all the case of N=2, 4, 8, and 12. The stable lumen size increases with increasing $N$.

In the experiment of epiblast, the rapid diffusion of dextran from the ECM side into the lumen suggested a large leak of the luminal fluid, while MDCK and pancreas did not show such evidence. This validates a low $\Delta p$ in epiblast and higher $\Delta p$ for MDCK and pancreas, and explains why lumens are formed in MDCK and pancreas at early stage and at each cell division thanks to higher $\Delta p$, while in epiblast microlumens cannot expand due to the smallness of $\Delta p$. It will take some time for the number of cells to increase sufficiently and for the apical surfaces of the cells to meet each other inside the organoid to nucleate a lumen via rosette structure.

*Footnote:

For N=12 case, since the same construction of equally segmented sphere is not possible, we consider regular dodecahedron with 12 faces as the shape of lumen and outer shape of organoid. In this configuration, each cell shape is a pentagonal pyramid whose tip is cut by a lumen with a self-similar pentagonal pyramid. In this case, each term in Eq. (20) is defined as

$$V_A = \frac{1}{2}(7\phi + 4)b^3, \qquad A_A = 3\sqrt{5(4\phi + 3)}b^2, \quad \gamma_L \sum_{i=1}^{n} A_L^i = 30\gamma_L \sqrt{\frac{1}{4}(3\phi^2 - 1)}\,(a^2 - b^2)$$

$k\sum_{j=1}^{N}\left(V_c^j - V_c^0\right)^2 = \frac{k}{12}\frac{1}{4}(7\phi+4)^2(a^3-b^3-a_0^3)^2$ , $k_E(V_E - V_E^0)^2 = k_E(g_l L^3 - \frac{1}{2}(7\phi+4)a^3 - g_l L_0^3)^2$ where $\phi = \frac{1+\sqrt{5}}{2}$, and a and b can be written by using the radius of circumscribed sphere of the lumen r, and that of organoid $R$ as $a = \frac{2R}{\sqrt{3}\phi}$, $b = \frac{2r}{\sqrt{3}\phi}$.
